# Juvenile reinstatement of TCF4 in Pitt-Hopkins syndrome model mice reveals a critical window for genetic intervention

**DOI:** 10.64898/2025.12.23.696277

**Authors:** Lucas M. James, Carlee A. Friar, Siyuan Liang, Eric B. Gao, Alain. C. Burette, Benjamin D. Philpot

**Affiliations:** UNC Neuroscience Center, University of North Carolina at Chapel Hill; Chapel Hill, NC 27599, USA; Department of Cell Biology and Physiology, University of North Carolina at Chapel Hill; Chapel Hill, NC 27599, USA; Carolina Institute for Developmental Disabilities, University of North Carolina at Chapel Hill; Campus Box #7255, Chapel Hill, NC 27599, USA

## Abstract

Pitt-Hopkins syndrome (PTHS) is a neurodevelopmental disorder caused by haploinsufficiency of *TCF4* which encodes transcription factor 4. As PTHS therapeutics advance toward clinical trials, identifying the optimal timing for treatment is crucial. Our previous research demonstrated that restoring TCF4 during embryonic or neonatal stages, corresponding to prenatal or neonatal periods in humans, improved phenotypes in a PTHS mouse model (Kim et al., 2022). However, PTHS diagnosis generally occurs much later, when infants fail to reach developmental milestones and undergo genetic testing. This raises an essential question: can genetic therapeutics initiated at more clinically relevant time points retain effectiveness? Here, we examined whether reinstating TCF4 in juvenile PTHS model mice could reverse behavioral phenotypes, simulating a gene therapy. Our findings indicate that this delayed intervention largely fails to correct most phenotypes, except for a measure of cognitive function. These results reveal phenotype-specific plasticity and underscore a narrow, early critical window for effective treatment in PTHS. Our study also identifies the hippocampus as a potential target for PTHS therapeutics and suggests that while some cognitive functions may still retain therapeutic plasticity, reversing most core PTHS symptoms may require intervention during the very early postnatal, or potentially prenatal periods, in humans.

## INTRODUCTION

Pitt-Hopkins syndrome (PTHS) is a neurodevelopmental disorder caused by haploinsufficiency of the *TCF4* gene which encodes the transcription factor 4 protein (TCF4) (Amiel et al., 2007; Brockschmidt et al., 2007; Christiane Zweier et al., 2007). Individuals with PTHS typically exhibit intellectual disability, lack of speech, motor delay, sleep disturbances, microcephaly, breathing disruption, hypotonia, gastrointestinal problems, and, in some cases, seizures (Goodspeed et al., 2018; Marangi et al., 2011; Zweier et al., 2008). Despite its high symptom burden, there is currently no disease-modifying therapeutic for PTHS, and clinical care is limited to symptom management (Goodspeed et al., 2018; Zollino et al., 2019). PTHS symptoms emerge early in life, with hypotonia and missed developmental milestones being key early indicators; however, a definitive diagnosis requires genetic testing (Marangi & Zollino, 2015; Whalen et al., 2012). Advances in access to clinical genetic testing have significantly reduced the average age of diagnosis of many neurodevelopmental disorders, however, PTHS is not yet included in newborn screening panels (Kernohan & Boycott, 2024; Zhang et al., 2024). This may change with the clinical success of a disease-modifying therapy.

While TCF4 biology remains poorly understood, it has been implicated in key neuronal processes during both early brain development and adulthood. TCF4 is most highly expressed during early development, with levels peaking perinatally in mice and humans (Rannals et al., 2016; Sirp et al., 2022). Additionally, TCF4 plays a role in neural progenitor proliferation, neuronal migration, and laminar organization in the developing forebrain (Li et al., 2019; Mesman et al., 2020; Page et al., 2018). Despite lower expression in the adult brain, TCF4 remains crucial for maintaining neuronal excitability, synaptic function, and plasticity throughout adulthood (Badowska et al., 2020; Brzózka et al., 2010; D’Rozario et al., 2016; Kennedy et al., 2016; Rannals et al., 2016). Beyond its essential roles in neurons, additional studies have linked TCF4 to the differentiation of oligodendrocyte progenitor cells and the development of oligodendrocytes and myelination (Bohlen et al., 2023; Furlanetto et al., 2025; Phan et al., 2020).

Although it has not been previously tested in PTHS models, early reinstatement of TCF4 during neurodevelopment is expected to provide the greatest therapeutic benefit similar to other neurodevelopment-related genes and their respective disorders. Therefore, preclinical studies have focused on modeling early intervention using both PTHS patient-derived brain organoids and mouse models (Kim et al., 2022; Papes et al., 2022). While these studies demonstrate the potential of reinstating TCF4 early in development, they most closely mirror an embryonic reinstatement of TCF4 in humans rather than a postnatal intervention, and as a result, may present an overly optimistic model of viable treatment strategies for PTHS that does not align with the typical timeline of diagnosis. Thus, there is a need to better define the optimal window for TCF4 reinstatement and identify appropriate outcome measures for such treatments.

Accordingly, we investigated whether restoring TCF4 expression during the juvenile period provides therapeutic benefits in a mouse model of PTHS. Using a conditional TCF4 reinstatement mouse previously developed and characterized by our lab, we modeled an idealized gene therapy reinstatement approach, in which all TCF4 isoforms were reinstated at appropriate levels in transduced neurons, allowing us to focus on assessing the potential of this later intervention. Together with our previous results, our findings suggest that perinatal reinstatement of TCF4 holds the greatest potential to treat behavioral symptoms of PTHS, with likely vastly diminishing therapeutic benefits upon juvenile reinstatement. However, our study also encouragingly suggests that some cognitive behaviors may still be responsive to TCF4 reinstatement later in life.

## RESULTS

### Retro-orbital injection of AAV-PHP.eB-hSyn1-Cre yields widespread biodistribution and rapid recombination

We previously demonstrated that delivering genetically-encoded Cre recombinase via adeno-associated virus (AAV) can reliably reinstate TCF4 expression in transduced neurons of the conditional reinstatement *Tcf4*-lox-stop-lox (*Tcf4*-LSL) mouse model, as Cre-mediated recombination removes a floxed transcriptional stop cassette downstream of exon 17 (Kim et al., 2020, 2022). Using this mouse model, we modeled an idealized “juvenile” onset TCF4 reinstatement gene therapy by injecting AAV-PHP.eB-hSyn1-Cre into the retro-orbital (RO) sinus of postnatal day 14 (P14) mice (Yardeni et al., 2011). This systemic delivery strategy confers widespread Cre biodistribution and recombination throughout the brain and in neurons, leveraging the ability of the AAV-PHP.eB capsid to traverse the blood-brain-barrier in C57BL/6J mice (Chan et al., 2017; Hordeaux et al., 2018).

We first established the biodistribution and time course over which injection of AAV-PHP.eB-hSyn1-Cre leads to recombination using Ai9 reporter mice (B6.Cg-*Gt(ROSA)26Sor^tm9(CAG-tdTomato)Hze^*/J) (Fig. 1A) (Madisen et al., 2010). Ai9 mice contain a loxP-STOP-loxP cassette similar to *Tcf4*-LSL mice, but Cre-mediated excision of the stop cassette in these mice induces expression of a tdTomato red fluorescent protein and thereby reports recombination efficiency with high fidelity and sensitivity (Madisen et al., 2010). We performed RO injections in P14 Ai9 mice, delivering 2.4e13 vg/kg of AAV-PHP.eB-hSyn1-Cre virus, and subsequently assessed tdTomato native immunofluorescence and Cre-recombinase antibody-mediated immunofluorescence in the brains of mice 4, 7, 10, and 17 days after injection (Fig. 1A). Delivery by RO injection yielded a widespread biodistribution of tdTomato and Cre proteins in neurons throughout the brain. Further, we detected tdTomato positive neurons as early as 4 days post-injection, with the greatest labeling occurring early in the cortex, hippocampus, thalamus, and midbrain (Fig. 1B). However, labeling increased in specific regions and peaked between 10 and 17 days following RO injection (Fig. 1B-E). The spatial distribution of tdTomato closely mirrored Cre biodistribution, demonstrating that the AAV-delivered Cre successfully induced recombination in transduced neurons (Fig. 1A-E).

**Figure 1.**
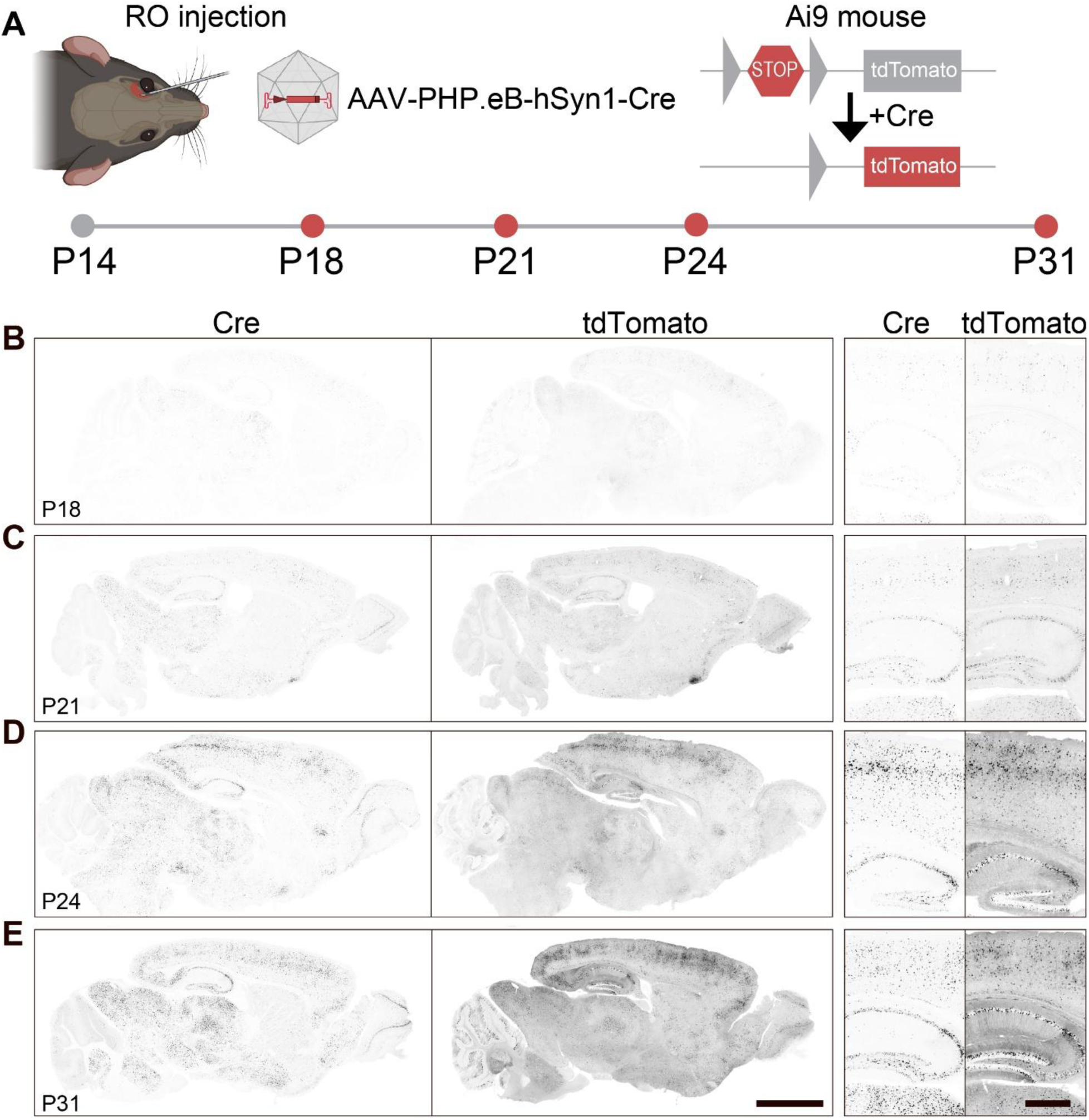
Injection of AAV-PHP.eB-hSyn1-Cre into the retro-orbital sinus yields widespread biodistribution and rapid recombination in the brain. **(A)** Schematic depicting injections of AAV-PHP.eB-hSyn1-Cre into the retro-orbital sinus (RO) of P14 Ai9 reporter mice and experimental time course of biodistribution assessment. **(B-E)** Representative Cre recombinase immunofluorescence and resulting tdTomato immunofluorescence in grayscale at the indicated ages. Scale bars, 2 mm (low-magnification) or 0.5 mm (high-magnification).

Importantly, we observed Cre expression in neurons throughout the cerebral cortex and the hippocampus, regions in which TCF4 expression is particularly high in mice, NHPs, and humans (Burette et al., 2024; Jung et al., 2018; Kim et al., 2020; Sirp et al., 2022) (Fig 1A-E). Neurons in the CA2 subregion of the hippocampus exhibited especially strong labeling, a region of previously documented preferential tropism by the AAV-PHP.eB capsid (Okamoto et al., 2023). Further, we observed strong Cre expression in subcortical structures, including the thalamus, hypothalamus, and striatum, areas where TCF4 is expressed in neurons, albeit at lower levels (Burette et al., 2024; Jung et al., 2018; Kim et al., 2020; Sirp et al., 2022). We also observed strong Cre labeling in the mitral cell layer of the olfactory bulb, the midbrain, and the hindbrain, although labeling in the cerebellum was primarily limited to the molecular and Purkinje cell layers rather than the granule cell layer.

Taken together, our Cre biodistribution demonstrates that RO injection of AAV-PHP.eB-hSyn1-Cre can readily achieve widespread delivery of Cre, and suggests that TCF4 reinstatement will follow a similar distribution in *Tcf4*-LSL mice. Additionally, the timing and pattern of tdTomato expression suggests that RO injection of AAV-PHP.eB-hSyn1-Cre at P14 in *Tcf4*-LSL mice will induce rapid recombination in transduced cells, with recombination reaching a plateau approximately two weeks following viral vector delivery.

### Reinstatement of TCF4 in juvenile PTHS model mice selectively improves spatial learning and memory

We next assessed the extent to which TCF4 reinstatement by AAV-PHP.eB-hSyn1-Cre delivery in juvenile mice mitigates behavioral deficits of *Tcf4*-LSL mice (Fig. 2A). To generate experimental litters, heterozygous male *Tcf4*-LSL mice were bred with wild-type (WT) females, producing WT and *Tcf4*-LSL littermates that received AAV-PHP.eB-hSyn1-Cre or AAV-PHP.eB-hSyn1-mCherry, as a negative control, via RO injection at P14. Before beginning behavioral assays, we verified the expression and biodistribution of the AAV-PHP.eB-hSyn1-Cre and AAV-PHP.eB-hSyn1-mCherry control vector at 2.4e13 vg/kg in *Tcf4*-LSL mice (Fig. 2B, C). We found that these vectors exhibited a similar distribution of Cre or mCherry, respectively, indicating that the viral vectors were appropriate for experimental comparisons.

**Figure 2.**
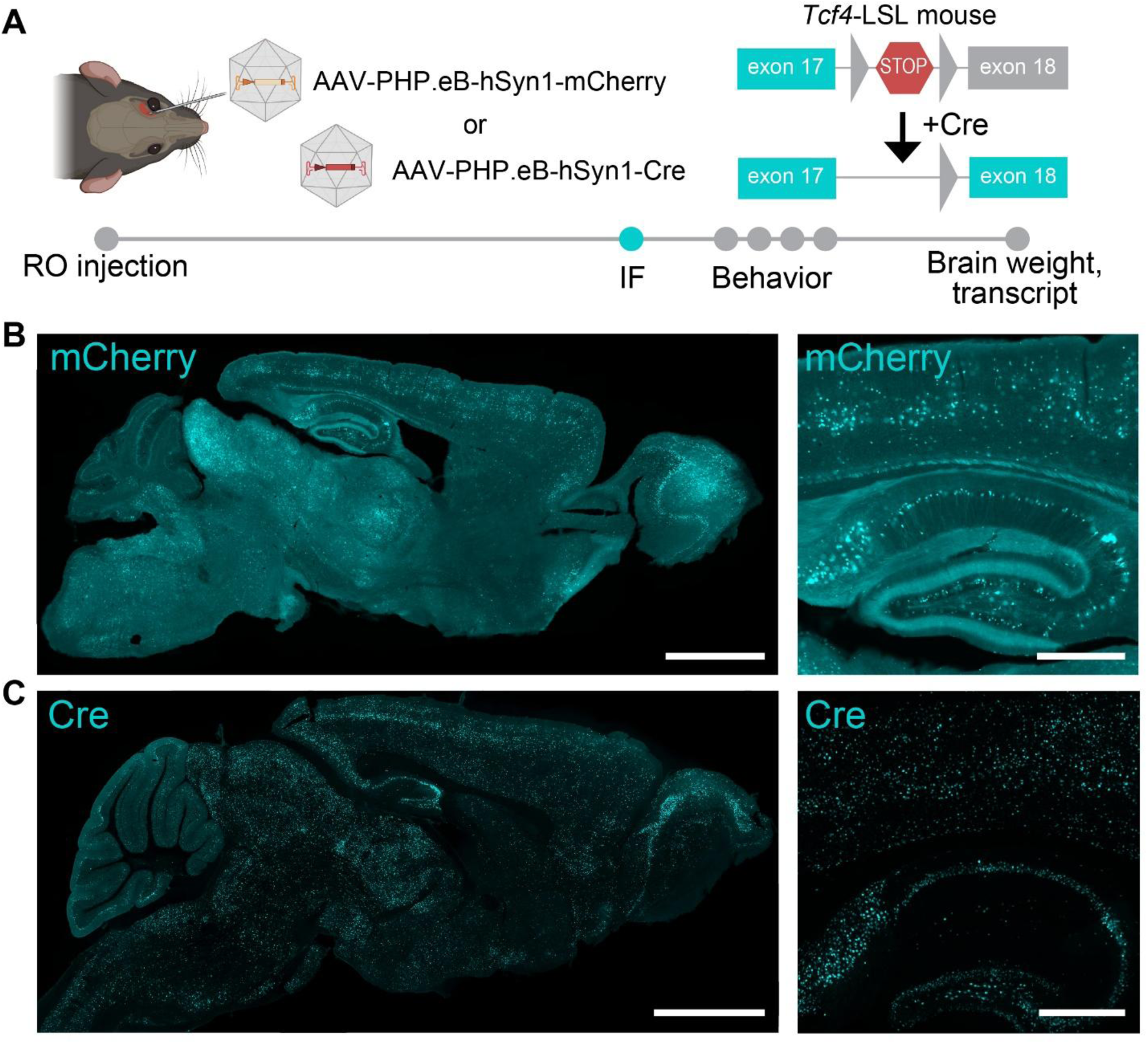
Representative biodistribution of AAV-hSyn1-mCherry and AAV-hSyn1-Cre in adult *Tcf4*-LSL mice. **(A)** Schematic depicting study paradigm including retro-orbital injection of AAV-PHP.eB-hSyn1-mCherry and AAV-PHP.eB-hSyn1-Cre in *Tcf4*-LSL mice, assessment of mCherry (native) or Cre (antibody-enhanced) immunofluorescence (IF) at postnatal day 60 and behavioral assessments and postmortem measures in mice separate from immunofluorescence. **(B,C)** Representative immunofluorescence for mCherry and Cre at postnatal day 60. Scale bars, 2 mm (low-magnification) or 0.5 mm (high-magnification).

Behavioral testing began when mice reached 9 weeks of age, using a behavioral battery previously used to phenotype PTHS model mice (Fig. 3A) (Kim et al., 2022).

**Figure 3.**
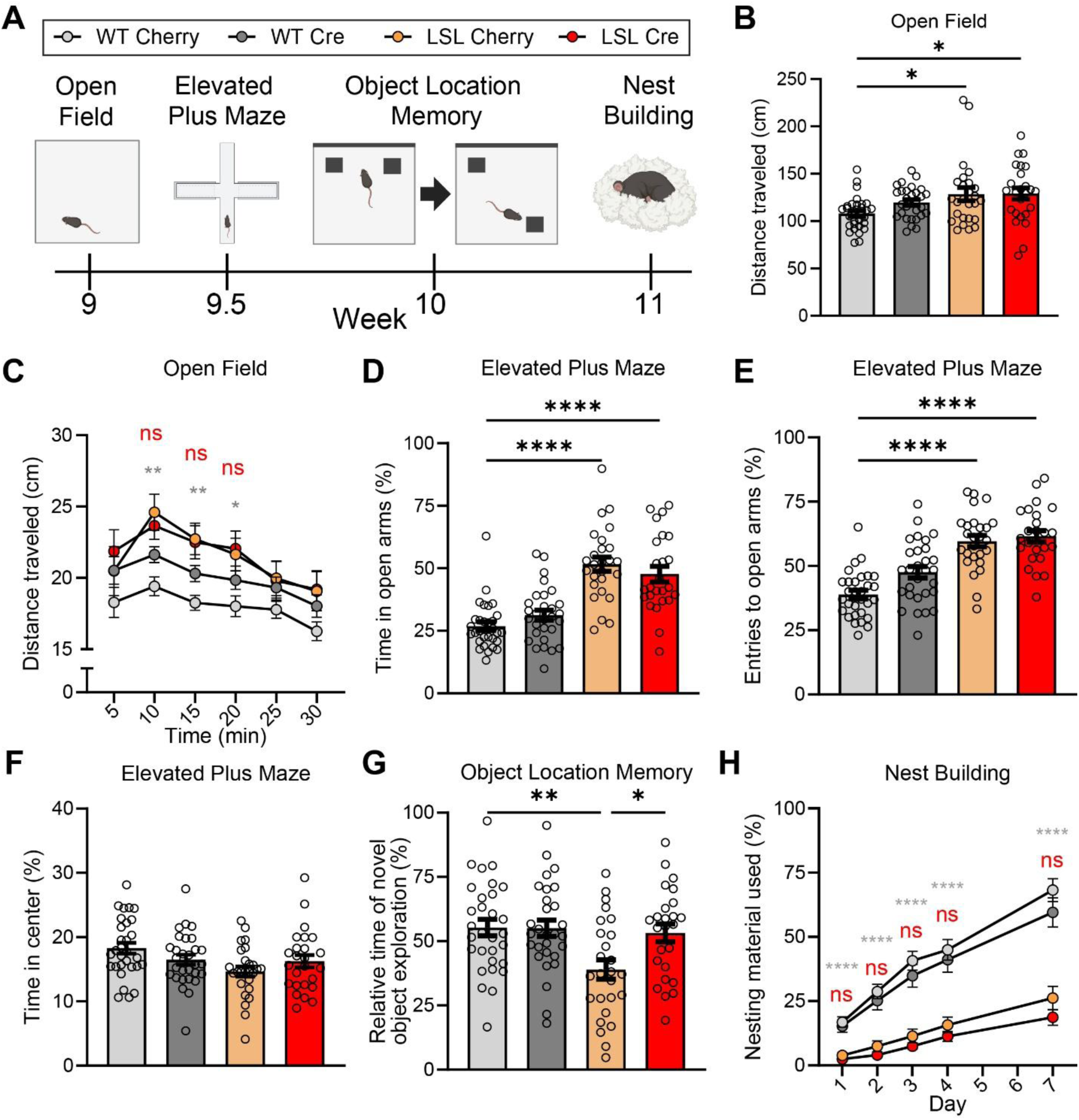
Behavioral recovery following juvenile reinstatement of TCF4 in *Tcf4*-LSL mice is limited to spatial memory. **(A)** Schematic of behavioral study timeline and behavioral assays in WT and *Tcf4*-LSL mice, following injection with AAV-hSyn1-mCherry or AAV-hSyn1-Cre. **(B-G)** Behavioral assessments, *n* = 25-30 mice per group. **(B)** Total distance traveled in the open field assay. Main effect of group by one-way ANOVA (*p* < 0.0001). **(C)** Distance traveled over time in the open field assay. Main effect of genotype (*p* < 0.0001) and time (*p* = 0.0085) by two-way ANOVA. **(D)** Time spent in and (**E**) entries to the open arms of the elevated plus maze. Main effect of group by one-way ANOVA (*p* < 0.0001, *p* < 0.0001, respectively). **(F)** Time spent in the center of the elevated plus maze. No effect by one-way ANOVA. **(G)** Relative novel object exploration time in the object location memory assay. Main effect of group by one-way ANOVA (*p* = 0.0024). **(H)** Percentage of nesting material used over time in the nest building assay, *n* = 23-26 mice per group. Main effect of genotype (*p* < 0.0001), trial (*p* < 0.0001), and interaction (*p* < 0.0001) by two-way ANOVA. Comparisons between WT Cherry - LSL Cherry (gray) and LSL Cre-LSL Cherry (red) are plotted. Data are represented as means ± SEM. Tukey’s multiple comparisons *post hoc* testing. * *p* < 0.05, ** *p* < 0.01, *** *p* < 0.001, **** *p* < 0.0001.

These experiments included both male and female mice and used littermate controls. Beginning with the open field assay, we found that mCherry-injected *Tcf4*-LSL mice exhibited increased distance traveled compared to their WT littermates, replicating previous findings; however, Cre-mediated reinstatement of *Tcf4* in juvenile *Tcf4*-LSL mice failed to mitigate this hyperactivity phenotype (Fig. 3B). Further, reinstatement of TCF4 in juvenile *Tcf4*-LSL mice did not improve habituation-dependent activity when distance traveled was evaluated across the duration of the 30-minute trial (Fig. 3C).

We next assessed performance in the elevated plus maze assay, a task which assesses approach-avoidance behavior of mice in the face of aversive height, light, and, thereby, perceived danger. PTHS model mice have been shown to exhibit increased entries into the open arms as well as increased time in the open arms of the elevated plus maze, suggesting reduced anxiety (Dennys et al., 2024; Kennedy et al., 2016; Kim et al., 2022; Thaxton et al., 2018). Similarly, we found that *Tcf4*-LSL mice receiving AAV-PHP.eB-hSyn1-mCherry spent more time in the open arms of the elevated plus maze and had an increased relative frequency of entries to the open arms compared to their similarly treated WT littermate controls (Fig. 3D, E) (Kim et al., 2022). However, unlike the recovery that we previously observed when neonatal (P1) *Tcf4*-LSL mice were treated with AAV-PHP.eB-hSyn1-Cre, Cre-mediated reinstatement of TCF4 in juvenile *Tcf4*-LSL mice failed to reduce time spent in the open arms of the elevated plus maze or entries to the open arms (Fig. 3D, E) (Kim et al., 2022). We found no differences in the time spent in the center of the elevated plus maze across groups (Fig. 3F).

Mouse models of PTHS also exhibit cognitive impairments as determined by performance in spatial learning and memory tasks (Kennedy et al., 2016; Kim et al., 2022; Thaxton et al., 2018). We assessed potential improvements in learning and memory phenotypes following juvenile TCF4 reinstatement using the object location memory assay. In this test, mice were habituated to an open field chamber over several days and then introduced to two identical objects in different locations. After 24 hours, mice were reintroduced to the same objects, but one object was moved to a new position, and we quantified the percentage of time spent exploring the relocated object relative to total object exploration time. PTHS mice typically spend a lower proportion of time exploring the object in a novel location than WT mice in this task, indicating impaired long-term spatial memory (Kennedy et al., 2016; Kim et al., 2022; Thaxton et al., 2018). We found that juvenile reinstatement of TCF4 improved performance in the object location memory task. Cre-treated *Tcf4*-LSL mice displayed significantly increased time exploring the object in the novel location compared to *Tcf4*-LSL mice injected with the negative control vector (Fig. 3G). Further, *Tcf4*-LSL mice injected with AAV-PHP.eB-hSyn1-Cre performed similarly to WT controls, suggesting that TCF4 reinstatement significantly improved long-term spatial memory to near WT levels (Fig. 3G).

Next, we assessed potential improvement in the nest building assay, in which PTHS model mice display a reduced ability to build a nest (Kim et al., 2022). Thought to inform an animal’s general well-being and neurologic state, the nest building assay measures an animal’s innate drive or ability to build a nest, a complex task of mouse daily living important for heat conservation and protection from predators (Deacon, 2006). We recapitulated the deficient nest building phenotype in AAV-PHP.eB-hSyn1-mCherry-treated *Tcf4*-LSL mice compared to WT controls; however, *Tcf4*-LSL mice exhibited no improvement in nest building activity following juvenile reinstatement of TCF4 compared to mCherry control vector-injected *Tcf4*-LSL mice (Fig. 3H).

PTHS model mice also exhibit both reduced body weight and brain weight (Kim et al., 2022; Thaxton et al., 2018). Therefore, we measured body weight weekly from the time of injection until the start of behavioral assays at approximately 9 weeks of age and assessed potential effects on body weight during development. As expected given the lack of improvement previously observed with neonatal TCF4 reinstatement (Kim et al., 2022), we found that juvenile reinstatement of TCF4 did not normalize body weight phenotypes in male or female mice compared to control *Tcf4*-LSL mice injected with the negative control vector (Fig. 4A, B). We also assessed potential effects on brain weight post-transcardial perfusion in a subset of mice from the study and found that juvenile TCF4 reinstatement also failed to improve brain weight phenotypes in *Tcf4*-LSL mice similar to our previous study (Fig. 4C).

**Figure 4.**
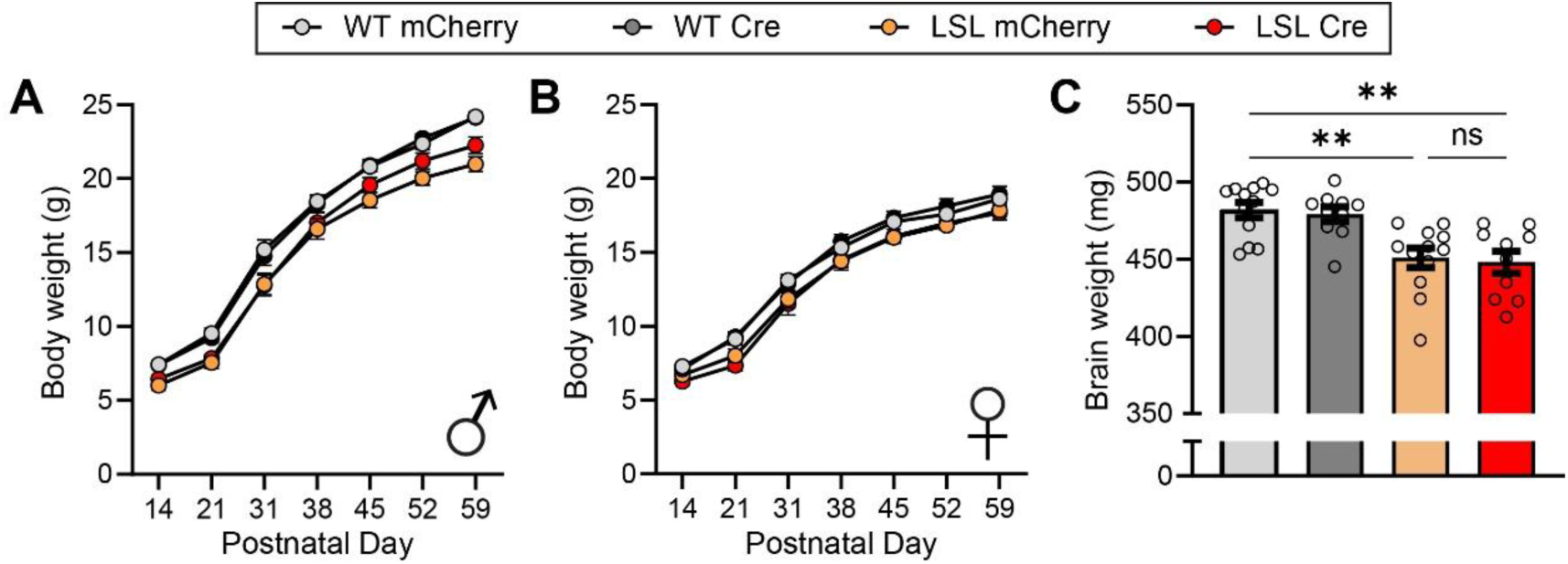
Juvenile reinstatement of TCF4 in *Tcf4*-LSL mice does not improve body weight or brain weight phenotypes. (A-B) Body weights of **(A)** male and (**B**) female WT and *Tcf4*-LSL mice following injection with AAV-hSyn1-mCherry or AAV-hSyn1-Cre, *n* = 12-16 mice per group. **(C)** Brain weights of WT and *Tcf4*-LSL mice following injection with AAV-hSyn1-mCherry or AAV-hSyn1-Cre, *n* = 10-12 mice per group. Comparisons between WT Cherry, LSL Cherry, and LSL Cre groups are plotted. Main effect of group by one-way ANOVA (*p* < 0.0001). Data are represented as means ± SEM. Tukey’s multiple comparisons *post hoc* testing. * *p* < 0.05, ** *p* < 0.01, *** *p* < 0.001, **** *p* < 0.0001.

### Assessment of TCF4 reinstatement and related impacts on transcriptional dysregulation in the hippocampus

Following the completion of behavioral experiments, we quantified the extent to which *Tcf4* levels were reinstated by AAV-PHP.eB-hSyn1-Cre. We found that RO injection of AAV-PHP.eB-hSyn1-Cre increased *Tcf4* levels by approximately 20% in the hippocampus compared to injection of AAV-PHP.eB-hSyn1-mCherry in a subset of mice from our behavioral study, confirming Cre-mediated reinstatement of *Tcf4* (Fig. 5A).

**Figure 5.**
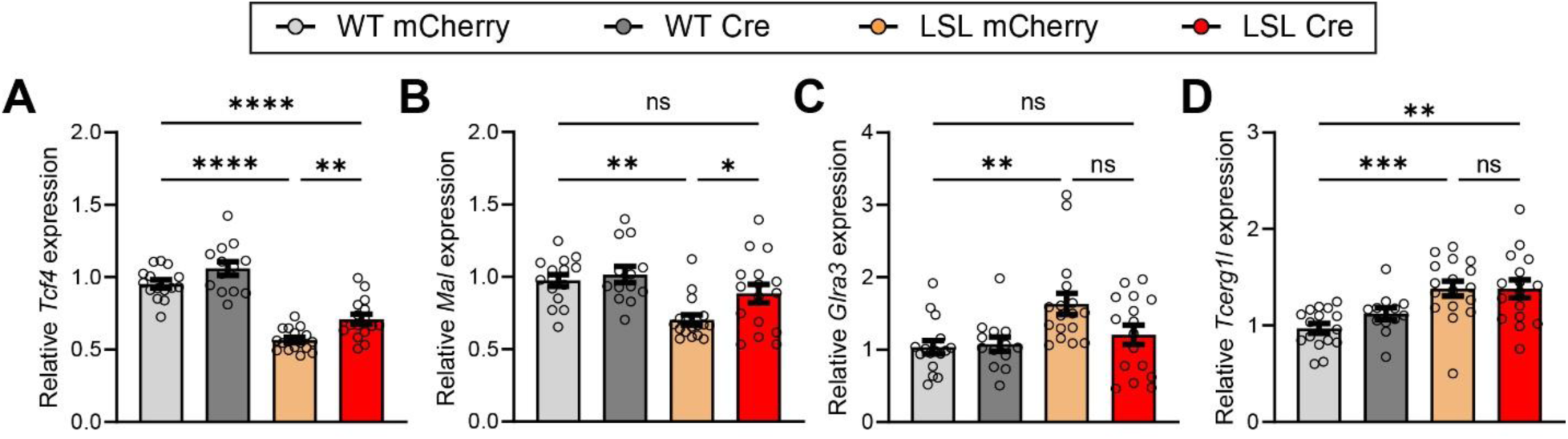
Juvenile reinstatement of TCF4 in *Tcf4*-LSL mice partially improves transcriptional dysregulation. (A-D) Relative expression of **(A)** *Tcf4*, **(B)** *Mal,* **(C)** *Glra3,* and **(D)** *Tcerg1l* in the hippocampus of WT and *Tcf4*-LSL mice following injection with AAV-hSyn1-mCherry or AAV-hSyn1-Cre, normalized to average of WT groups. Comparisons between WT Cherry, LSL Cherry, and LSL Cre groups are plotted, *n* = 13-17 mice per group. Main effect of group by one-way ANOVA (*p* < 0.0001, *p* = 0.0002, *p* = 0.0033, *p* = 0.0002 for A-D, respectively). Data represent means ± SEM. Tukey’s multiple comparisons *post hoc* testing. * *p* < 0.05, ** *p* < 0.01, *** *p* < 0.001, **** *p* < 0.0001.

Notably, this virus-mediated increase in *Tcf4* levels following P14 RO AAV-PHP.eB-hSyn1-Cre delivery is comparable to what we previously achieved following P1 ICV delivery, suggesting that the more modest phenotypic recovery with juvenile treatment resulted from reinstatement later in development, rather a difference in transduction efficiency (Kim et al., 2022).

Finally, we assessed potential impacts on transcriptional dysregulation observed in models of *TCF4* haploinsufficiency. As TCF4 is a transcription factor, several studies have identified genes that are dysregulated by heterozygous loss of *Tcf4*, though it remains unclear whether these genes are directly regulated by TCF4 or indirectly dysregulated as a downstream consequence of haploinsufficiency (Forrest et al., 2018; Kennedy et al., 2016; Li et al., 2019; Phan et al., 2020). Nonetheless, these dysregulated genes can serve as a readout for transcriptional normalization following TCF4 reinstatement. Accordingly, we used RT-qPCR to assess transcript levels of some of these candidate genes in the hippocampus following TCF4 reinstatement. We found that *MaI* levels were significantly improved by juvenile reinstatement of TCF4, while levels of *Glra3* and *Tcerg1l* were not significantly improved (Fig. 5B-D). This suggests that while some transcriptional dysregulation can be improved by TCF4 reinstatement in juvenile mice, the potential for improvements may be limited.

## DISCUSSION

The convergence of widespread clinical genetic testing with advances in molecular therapeutics signals a pressing need to identify optimal treatment windows for genetic disorders, especially for neurodevelopmental disorders like PTHS. We previously found that embryonic or neonatal reinstatement of TCF4 in PTHS model mice could improve behavioral and transcriptional dysregulation phenotypes, painting an optimistic picture for PTHS gene therapy approaches (Kim et al., 2022). This was bolstered by findings that increasing TCF4 expression in PTHS patient-derived brain organoids could overcome deficits in electrophysiological, cytoarchitectural, and transcriptional dysregulation phenotypes (Papes et al., 2022). However, these studies modeled very early, likely embryonic, timepoints of intervention in humans that do not reflect clinical reality. Here, we simulated a TCF4 upregulation-based gene therapy, administered on a timeline aligned with postnatal diagnosis and intervention, to establish the extent to which reinstatement of TCF4 later in development might be therapeutic. In stark contrast with embryonic and neonatal intervention in mice, we found that most behavioral phenotypes were not reversed following intervention at this later timepoint, except a measure of specific cognitive abilities (summarized in Table 1). These findings temper expectations for postnatal TCF4 reinstatement therapies, including gene-addition strategies now approaching clinical trials, yet suggest that certain cognitive domains may remain amenable to intervention in young children.

**Table 1.** Summary of mouse behavioral and anatomical phenotypes with demonstrated recovery (R) or no recovery (NR) following TCF4 reinstatement at various stages of development from Kim et al., 2022 and the present study.

| Phenotype | Timing of intervention |  |  |
| --- | --- | --- | --- |
|  | Embryonic | P1 | P14 |
| Object Location Memory | R | R | R |
| Elevated Plus Maze | R | R | NR |
| Nest Building | R | R | NR |
| Open Field | R | R | NR |
| Brain Weight | R | NR | NR |
| Body Weight | R | NR | NR |

Our goal was to model a realistic gene therapy approach, simulating intervention at a time after diagnosis. Following P14 delivery of AAV-PHP.eB-hSyn1-Cre, transgene expression increased over approximately two weeks – a developmental period in mice marked by the rapid transition from just after eye opening and early sensory maturation to near-complete achievement of major neurodevelopmental milestones (Kronman et al., 2024; Zalewska, 2019). However, differences in neurodevelopmental timing across species prevent direct and precise extrapolation to humans (Cottam et al., 2024; Wallace & Pollen, 2024; Workman et al., 2013). By analogy, this intervention roughly corresponds to reinstatement at the toddler / early childhood developmental stage in humans, representing a feasible yet conservative intervention window for individuals with PTHS.

The improvement we observed in a cognitive task (object location memory) highlights the promising possibility that cognitive improvements might be achievable in individuals with PTHS through effective TCF4 reinstatement. Further, the selective rescue of object location memory is particularly mechanistically interesting. This task depends on the hippocampus, a region that remains plastic and in which TCF4 expression is high throughout life (Moser et al., 1993; Moser et al., 1995; Nakazawa et al., 2004). While TCF4 levels decline sharply during the perinatal period in many brain regions, the adult hippocampus retains significant TCF4 expression in mice, non-human primates, and humans, suggesting that TCF4 continues to play an important role in this part of the brain (Burette et al., 2024; Jung et al., 2018; Kim et al., 2020; Sirp et al., 2022). Additionally, hippocampal neurons retain substantial synaptic plasticity throughout life, potentially positioning the hippocampus as a key target for therapeutic recovery of cellular and synaptic function (Bittner et al., 2017; Nakazawa et al., 2004).

TCF4’s known role in regulating dendritic spine density and shape supports the potential for synaptic remodeling after reinstatement, since restoring TCF4 function could promote activity-dependent spine plasticity that underlies learning and memory (Badowska et al., 2020; Crux et al., 2018; Li et al., 2019; Sarkar et al., 2021). Given the strong TCF4 reinstatement achieved in the hippocampus and the resulting improvement in hippocampus-dependent spatial memory, our findings suggest targeting TCF4 delivery specifically to this region might be a feasible therapeutic approach if broad CNS reinstatement of TCF4 is not possible.

Our findings must be interpreted with several important caveats. Despite our observation that a cognitive task and at least some transcriptional signatures can be ameliorated with our late-onset TCF4 reinstatement approach, several arguments can be made that we may be overestimating the benefits of a gene therapy for PTHS. First, we employed the PHP.eB capsid, which exhibits substantially greater efficacy in mice than readily available capsids in primates (Bey et al., 2020; Chan et al., 2017). Our viral biodistribution thus exceeds what could be likely achieved with AAV9 in humans, potentially overestimating the benefits of postnatal TCF4 reinstatement in humans (Bailey et al., 2020; Foust et al., 2009; Hinderer et al., 2014). Second, Cre-mediated excision of the STOP cassette restores native expression of TCF4 in transduced neurons, mitigating risks inherent to gene therapy where under-expression might fail to correct cellular deficits or over-expression of TCF4 could produce cellular toxicity and network dysfunction (Badowska et al., 2020; Brzózka et al., 2010). Third, our reinstatement strategy also recovers the endogenous expression ratio of TCF4 isoforms, which would not be feasible with traditional gene delivery approaches, as AAV packaging limits make delivering multiple isoforms of TCF4 challenging. At least 18 TCF4 isoforms are expressed in humans, and the evolutionarily-conserved complexity of *TCF4* splicing and developmental regulation suggests these isoforms may serve distinct biological functions (Sirp et al., 2022). Therefore, our findings may be most relevant for *TCF4* upregulation therapeutic approaches and may especially overestimate the performance of gene addition strategies.

Conversely, several factors suggest we may be underestimating therapeutic potential. Next-generation capsids demonstrating great promise for broad neuronal transduction are actively under development (Goertsen et al., 2022; Huang et al., 2024; Moyer et al., 2025), and dramatically improved biodistribution might yield broader therapeutic benefits than we observed. Additionally, we selectively reinstated TCF4 in neurons using the Syn1 promoter (Kügler et al., 2003). While TCF4 is predominantly neuronal in both mice and primates, TCF4 is also expressed in glia (Burette et al., 2024; Kim et al., 2020), and emerging evidence highlights the importance of oligodendrocyte TCF4 expression (Bohlen et al., 2023; Phan et al., 2020). Notably, neuronal TCF4 reinstatement partially normalized *Mal* expression, a transcript nearly exclusive to oligodendrocytes in the brain (Schaeren-Wiemers et al., 1995), suggesting non-cell autonomous therapeutic effects following neuronal TCF4 reinstatement. Finally, the inherent limitations of assessing cognition in mice raise the possibility that postnatal TCF4 reinstatement in humans could confer cognitive benefits broader than those which can be appreciated in mice.

In summary, our study provides the first evidence that TCF4 reinstatement early in development is necessary to maximize therapeutic benefit in PTHS, adding PTHS to the growing list of neurodevelopmental disorders in which mouse models have shown such a critical period which includes Angelman syndrome, Phelan-McDermid syndrome*, GRIN1*-related disorders, and *SYNGAP1-*related cognitive impairment (Aceti et al., 2015; Clement et al., 2012; Mei et al., 2016; Mielnik et al., 2021; Silva-Santos et al., 2015). Our findings that TCF4 reinstatement provides little benefit in juvenile mice beyond a measure of cognitive function could serve as an important bellwether for upcoming AAV-based TCF4 addition clinical trials and gene therapies in which (1) the biodistribution of TCF4 reinstatement may be limited, (2) only one or two isoforms may be expressed given viral packaging limits, and (3) the under-or over-expression of TCF4 in individual cells present additional challenges. In addition to developing technologies that overcome some of these limitations, future gene therapy efforts in PTHS should explore approaches to upregulate the intact *TCF4* allele, such as CRISPRa approaches, particularly in manners that may be tunable.

## MATERIALS AND METHODS

### Animal use

All animal procedures performed in this study were approved by the Institutional Animal Care and Use Committee at the University of North Carolina at Chapel Hill. Mice were raised and maintained on a 12:12 light-dark cycle with *ad libitum* access to food and water. *Tcf4*-LSL mice were previously generated and are maintained on a C57BL/6J background (Kim et al., 2020). Mice were genotyped using primers described in Supplemental Table 1.

### Virus production

Plasmids for virus production were obtained from Addgene (pAAV-hSyn-mCherry, #114472; pENN.AAV.hSyn.Cre.hGH, #10555). AAV serotype AAV-PHP.eB was produced using polyethyleneimine triple transfection and purified by three rounds of CsCl density gradient centrifugation. Purified vector was exchanged into 0.1 mM phosphate-buffered saline (PBS) containing 5% D-Sorbitol and 350 mM NaCl.

### Retro-orbital injections

Immediately prior to injections, postnatal day 14 (P14) mice were weighed for vector dosage calculations. Mice were anaesthetized by inhalation of isoflurane (Piramal) at 2-2.5% in O_2_ for 3-4 minutes in an airtight chamber. Lack of toe pinch reflex was verified and then mice were placed on their left side in the lateral position. A 50 μl syringe fitted with a 32-gauge, 0.4-inch-long sterile syringe needle point style 4 (Hamilton, #7655–01) was used for injections into the retro-orbital sinus. Virus was diluted immediately prior to each injection in vehicle solution (0.1 mM PBS, 5% D-Sorbitol, and 350 mM NaCl). Mice were injected with 2.4 x 10^13^ vector genomes/kg in a total volume of 35 μl and recovered from anesthesia on a heating pad. Body weight was monitored following injection to monitor for toxicity.

### Tissue collection and preparation

For histology, mice received intraperitoneal injections of Euthasol (100 mg/kg) for deep anesthesia and were perfused (intracardially) with PBS followed by 4% paraformaldehyde in PBS, pH 7.4. Following perfusion, brains were extracted, postfixed in 4% paraformaldehyde in PBS for 24 hours, and stored in 30% sucrose in PBS at 4°C. For molecular biology, mice were rapidly euthanized, and brains were extracted in cold PBS, pH 7.4. Brains were split along the midline, and hippocampus was dissected, snap-frozen, and stored at-80°C until processing.

### Immunohistochemistry

Perfused brains were cut along the midline, and one hemisphere was sectioned in sagittal orientation at a thickness of 40 μm using a sliding microtome. Sections were stored at-20°C in cryoprotective solution (45% PBS, 30% ethylene glycol, 25% glycerol) until staining. Free-floating sections were washed several times in PBS and PBS with 0.1% Triton X-100 (PBST) before a 30-minute blocking step in 10% Fetal Bovine Serum (FBS) in PBST at room temperature (RT). Sections were then incubated with Cre-recombinase primary antibody (Synaptic Systems, #257003, 1:500) in 10% FBS in PBST for 24 hours at RT. Sections were then washed in PBST and incubated for 24 hours in 10% FBS in PBST with secondary antibody (Jackson Immuno, #711605152, 1:400) and DAPI (ThermoFisher, #D21490, 1 µg/ml). The next day, sections were washed with PBST and PBS and mounted on gelatin-covered glass slides and Vectashield Plus Antifade Mounting Medium (Vector Labs, #H-1900-10) was used for coverslipping. Slides were imaged on a Leica STELLARIS 8 FALCON or Olympus VS200.

### Mouse behavioral assays

All experimental animals used in the behavioral study were produced by breeding a WT C57BL/6J dam with a C57BL/6J heterozygous *Tcf4*-LSL sire. Behavioral assays were performed in presumed order of least to most stressful, during the light cycle, and with at least 2-3 days in between assays. Sample sizes were determined by effect sizes and power calculations from previous studies in PTHS model mice. Male and female mice were run separately, and sex was not considered a biological variable, except for body weight, as both male and female mice display behavioral phenotypes. Sex-matched littermates were used for all experiments, and treatment assignment was random.

Equipment was cleaned in between animals using 10% ethanol and dried before beginning of the next trial.

Open Field: Mice were placed in an arena (40×40×30 cm) inside a sound-attenuating box equipped with ceiling-mounted lights. Animals were allowed to freely explore for 30 minutes. Distance traveled was measured and analyzed in 5-minute bins and the total duration of the trial using EthoVision XT 15.0.

Elevated Plus Maze: The elevated plus maze was elevated 50 cm above the floor and contains two open and two closed arms, with arms 20 cm in length, 8 cm in width, and walls 20 cm in height for the closed arms. Mice were placed on the center section and movements were recorded with a video camera. After 5 minutes of free exploration, mice were removed and time in and entries to the open and closed arms were quantified using EthoVision XT 15.0.

Object Location Memory: Mice were habituated to an open field arena contained inside a sound-attenuating box equipped with ceiling-mounted lights for 5 minutes each day for 3 days. The arena contained a visual cue on one side and did not contain any objects.

On the fourth day, mice were placed in the chamber with two identical objects and allowed to freely explore for 10 minutes. On the fifth day, mice were placed in the same chamber but one of the objects was moved to a novel location. Mice were allowed to explore the chamber for 5 minutes. Day 5 was video recorded and the interaction time of a mouse with each object was measured using EthoVision XT 15.0 and the percentage of time spent exploring the object in a novel position of total object exploration time was calculated.

Nest Building: Mice were single housed for 3 days prior to the start of the nest building assay and throughout its duration. On day 0, home nestlet was removed and extra thick filter paper (Bio-Rad, #1703966) was put in its place. Each cage received a total mass of 10.5-11.5 g of filter paper that was cut into 8 evenly sized rectangles. Starting on day 1, the unincorporated nestlet was weighed each day for 4 consecutive days and again on Day 7. The percentage of used nestlet was calculated by mass. Note, 13 animals began and did not finish the assay due to disruption to their home cage environment during the assay.

### RNA extractions and RT-qPCR

Following microdissection, hippocampal samples were flash frozen and stored at-80°C. Tissue samples were lysed and homogenized using a Tissue Tearor (Model 985–370) in RLT lysis buffer (Qiagen, #79216) with 1% β-mercaptoethanol. RNA was extracted according to manufacturer’s protocols using the RNeasy kit (Qiagen, #74106). cDNA was synthesized from 500 ng of RNA using qScript cDNA Supermix (Quantabio, #101414-106). PowerUp SYBR Green Master Mix (Applied Biosystems, #A25742) was used for qPCR using primers indicated in Supplemental Table 1 on a QuantStudio 5 Real-Time PCR system (Applied Biosystems). Relative expression was normalized to *Eif4a2* expression using the comparative CT method.

### Statistics and reproducibility

Statistical tests used are indicated in each figure legend by experiment and were calculated in GraphPad Prism 10. Experiments with more than two groups and one varying factor were analyzed by one-way analysis of variance (ANOVA) followed by Tukey’s multiple comparisons tests. Experiments with more than two groups and two varying factors were analyzed by two-way ANOVA with Tukey’s multiple comparisons *post hoc* tests. Differences were considered statistically significant if *p* < 0.05.

## Acknowledgements and funding

This research was supported by NIH NINDS grants R01NS114086, R01NS129914 and R01NS131615, the Pitt-Hopkins Research Foundation, and a donation from the Israeli Pitt-Hopkins Association to BDP, as well as a Royster Fellowship to LMJ. Microscopy was performed at the UNC Hooker Imaging Core Facility, supported in part by P30 CA016086 Cancer Center Core Support Grant to the UNC Lineberger Comprehensive Cancer Center, and NIH grant 1S10OD030300. Viral vectors were produced at the UNC BRAIN Initiative Viral Vector Core, supported by 5U24NS124025.

## Author contributions

Conceptualization: LMJ, BDP Data curation: LMJ, CAF, SL, EBG

Formal analysis: LMJ, CAF, SL, EBG Funding acquisition: LMJ, BDP Investigation: LMJ, CAF, SL, EBG, ACB

Methodology: LMJ, CAF, SL, EBG, ACB Project administration: LMJ Visualization: LMJ

Supervision: BDP

Writing – original draft: LMJ

Writing – review & editing: LMJ, CAF, SL, EBG, ACB, BDP

## Data and materials availability

Individual data points are represented in each figure and numerical data will be made publicly available in the final publication.

## SUPPLEMENTARY INFORMATION

**Supplementary Table 1.**
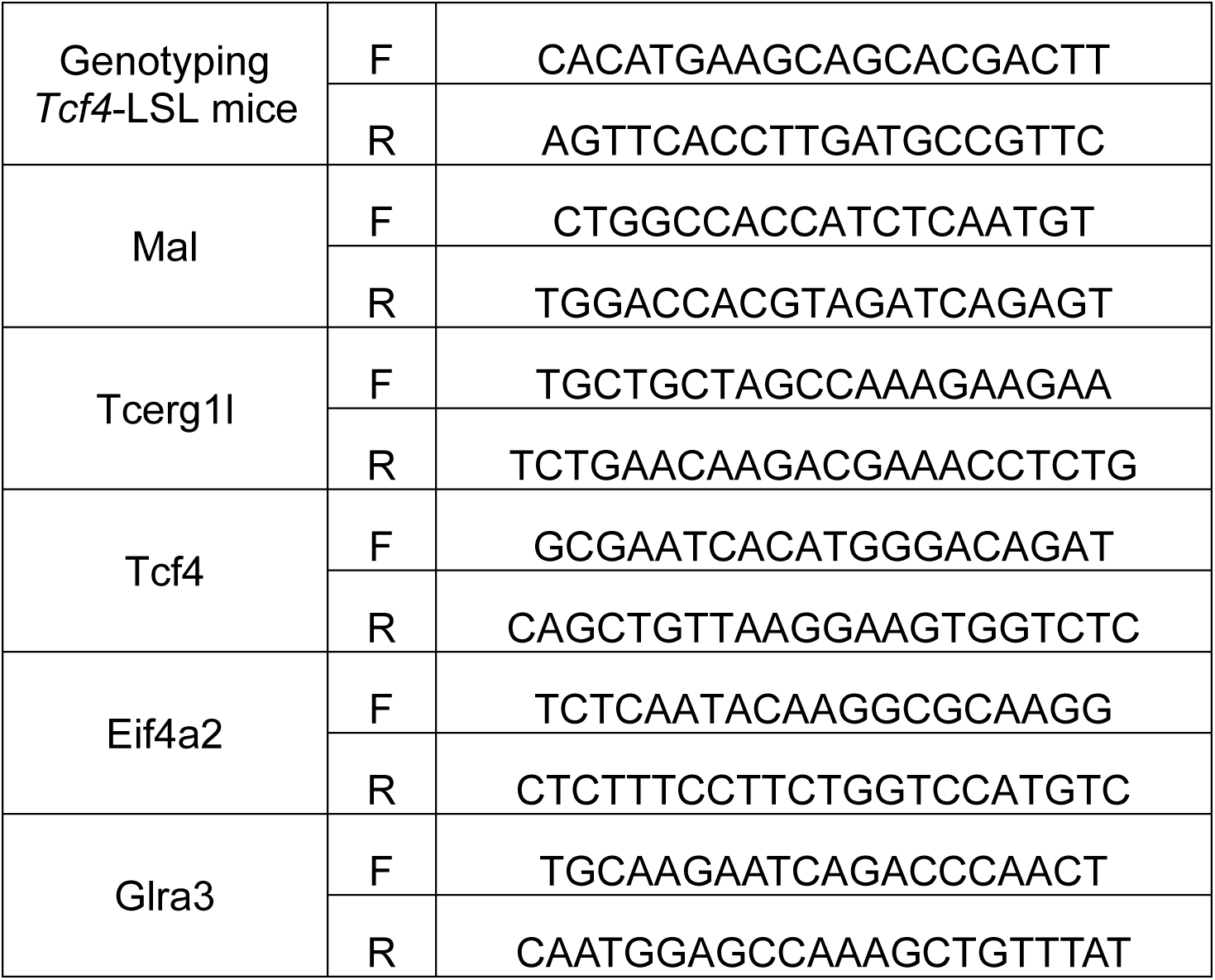
Primers used throughout study.

## REFERENCES

Aceti, M., Creson, T. K., Vaissiere, T., Rojas, C., Huang, W.-C., Wang, Y.-X., Petralia, R. S., Page, D. T., Miller, C. A., & Rumbaugh, G. (2015). Syngap1 haploinsufficiency damages a postnatal critical period of pyramidal cell structural maturation linked to cortical circuit assembly. Biological Psychiatry, 77(9), 805–815. 10.1016/j.biopsych.2014.08.001

Amiel, J., Rio, M., de Pontual, L., Redon, R., Malan, V., Boddaert, N., Plouin, P., Carter, N. P., Lyonnet, S., Munnich, A., & Colleaux, L. (2007). Mutations in TCF4, encoding a class I basic helix-loop-helix transcription factor, are responsible for Pitt-Hopkins syndrome, a severe epileptic encephalopathy associated with autonomic dysfunction. American Journal of Human Genetics, 80(5), 988–993. 10.1086/515582

Badowska, D. M., Brzózka, M. M., Kannaiyan, N., Thomas, C., Dibaj, P., Chowdhury, A., Steffens, H., Turck, C. W., Falkai, P., Schmitt, A., Papiol, S., Scheuss, V., Willig, K. I., Martins-de-Souza, D., Rhee, J. S., Malzahn, D., & Rossner, M. J. (2020). Modulation of cognition and neuronal plasticity in gain-and loss-of-function mouse models of the schizophrenia risk gene Tcf4. Translational Psychiatry, 10(1), 343. 10.1038/s41398-020-01026-7

Bailey, R. M., Rozenberg, A., & Gray, S. J. (2020). Comparison of high-dose intracisterna magna and lumbar puncture intrathecal delivery of AAV9 in mice to treat neuropathies. Brain Research, 1739, 146832. 10.1016/j.brainres.2020.146832

Bey, K., Deniaud, J., Dubreil, L., Joussemet, B., Cristini, J., Ciron, C., Hordeaux, J., Le Boulc’h, M., Marche, K., Maquigneau, M., Guilbaud, M., Moreau, R., Larcher, T., Deschamps, J.-Y., Fusellier, M., Blouin, V., Sevin, C., Cartier, N., Adjali, O.,…Colle, M.-A. (2020). Intra-CSF AAV9 and AAVrh10 Administration in Nonhuman Primates: Promising Routes and Vectors for Which Neurological Diseases? Molecular Therapy. Methods & Clinical Development, 17, 771–784. 10.1016/j.omtm.2020.04.001

Bittner, K. C., Milstein, A. D., Grienberger, C., Romani, S., & Magee, J. C. (2017). Behavioral time scale synaptic plasticity underlies CA1 place fields. Science, 357(6355), 1033–1036. 10.1126/science.aan3846

Bohlen, J. F., Cleary, C. M., Das, D., Sripathy, S. R., Sadowski, N., Shim, G., Kenney, R. F., Buchler, I. P., Banerji, T., Scanlan, T. S., Mulkey, D. K., & Maher, B. J. (2023). Promyelinating drugs promote functional recovery in an autism spectrum disorder mouse model of Pitt-Hopkins syndrome. Brain: A Journal of Neurology, 146(8), 3331– 3346. 10.1093/brain/awad057

Brockschmidt, A., Todt, U., Ryu, S., Hoischen, A., Landwehr, C., Birnbaum, S., Frenck, W., Radlwimmer, B., Lichter, P., Engels, H., Driever, W., Kubisch, C., & Weber, R. G. (2007). Severe mental retardation with breathing abnormalities (Pitt-Hopkins syndrome) is caused by haploinsufficiency of the neuronal bHLH transcription factor TCF4. Human Molecular Genetics, 16(12), 1488–1494. 10.1093/hmg/ddm099

Brzózka, M. M., Radyushkin, K., Wichert, S. P., Ehrenreich, H., & Rossner, M. J. (2010). Cognitive and sensorimotor gating impairments in transgenic mice overexpressing the schizophrenia susceptibility gene Tcf4 in the brain. Biological Psychiatry, 68(1), 33–40. 10.1016/j.biopsych.2010.03.015

Burette, A. C., Vihma, H., Smith, A. L., Ozarkar, S. S., Bennett, J., Amaral, D. G., & Philpot, B. D. (2024). Transcription factor 4 expression in the developing non-human primate brain: a comparative analysis with the mouse brain. Frontiers in Neuroanatomy, 18, 1478689. 10.3389/fnana.2024.1478689

Chan, K. Y., Jang, M. J., Yoo, B. B., Greenbaum, A., Ravi, N., Wu, W.-L., Sánchez-Guardado, L., Lois, C., Mazmanian, S. K., Deverman, B. E., & Gradinaru, V. (2017). Engineered AAVs for efficient noninvasive gene delivery to the central and peripheral nervous systems. Nature Neuroscience, 20(8), 1172–1179. 10.1038/nn.4593

Clement, J. P., Aceti, M., Creson, T. K., Ozkan, E. D., Shi, Y., Reish, N. J., Almonte, A. G., Miller, B. H., Wiltgen, B. J., Miller, C. A., Xu, X., & Rumbaugh, G. (2012). Pathogenic SYNGAP1 mutations impair cognitive development by disrupting maturation of dendritic spine synapses. Cell, 151(4), 709–723. 10.1016/j.cell.2012.08.045

Cottam, N. C., Ofori, K., Bryant, M., Rogge, J. R., Hekmatyar, K., Sun, J., & Charvet, C. J. (2024). From circuits to lifespan: translating mouse and human timelines with neuroimaging based tractography. BioRxiv. 10.1101/2024.07.28.605528

Crux, S., Herms, J., & Dorostkar, M. M. (2018). Tcf4 regulates dendritic spine density and morphology in the adult brain. Plos One, 13(6), e0199359. 10.1371/journal.pone.0199359

D’Rozario, M., Zhang, T., Waddell, E. A., Zhang, Y., Sahin, C., Sharoni, M., Hu, T., Nayal, M., Kutty, K., Liebl, F., Hu, W., & Marenda, D. R. (2016). Type I bHLH Proteins Daughterless and Tcf4 Restrict Neurite Branching and Synapse Formation by Repressing Neurexin in Postmitotic Neurons. Cell Reports, 15(2), 386–397. 10.1016/j.celrep.2016.03.034

Deacon, R. M. J. (2006). Assessing nest building in mice. Nature Protocols, 1(3), 1117– 1119. 10.1038/nprot.2006.170

Dennys, C. N., Vermudez, S. A. D., Deacon, R. J. M., Sierra-Delgado, J. A., Rich, K., Zhang, X., Buch, A., Weiss, K., Moxley, Y., Rajpal, H., Espinoza, F. D., Powers, S., Ávila, A. S., Gogliotti, R. G., Cogram, P., Niswender, C. M., & Meyer, K. C. (2024). MeCP2 gene therapy ameliorates disease phenotype in mouse model for Pitt Hopkins syndrome. Neurotherapeutics, 21(5), e00376. 10.1016/j.neurot.2024.e00376

Forrest, M. P., Hill, M. J., Kavanagh, D. H., Tansey, K. E., Waite, A. J., & Blake, D. J. (2018). The psychiatric risk gene transcription factor 4 (TCF4) regulates neurodevelopmental pathways associated with schizophrenia, autism, and intellectual disability. Schizophrenia Bulletin, 44(5), 1100–1110. 10.1093/schbul/sbx164

Foust, K. D., Nurre, E., Montgomery, C. L., Hernandez, A., Chan, C. M., & Kaspar, B. K. (2009). Intravascular AAV9 preferentially targets neonatal neurons and adult astrocytes. Nature Biotechnology, 27(1), 59–65. 10.1038/nbt.1515

Furlanetto, F., Flegel, N., Kremp, M., Spear, C., Fröb, F., Alfonsetti, M., Bohl, B., Krumbiegel, M., Turan, S., Reis, A., Lie, D. C., Winkler, J., Falk, S., Wegner, M., & Karow, M. (2025). A novel human organoid model system reveals requirement of TCF4 for oligodendroglial differentiation. Life Science Alliance, 8(6). 10.26508/lsa.202403102

Goertsen, D., Flytzanis, N. C., Goeden, N., Chuapoco, M. R., Cummins, A., Chen, Y., Fan, Y., Zhang, Q., Sharma, J., Duan, Y., Wang, L., Feng, G., Chen, Y., Ip, N. Y., Pickel, J., & Gradinaru, V. (2022). AAV capsid variants with brain-wide transgene expression and decreased liver targeting after intravenous delivery in mouse and marmoset. Nature Neuroscience, 25(1), 106–115. 10.1038/s41593-021-00969-4

Goodspeed, K., Newsom, C., Morris, M. A., Powell, C., Evans, P., & Golla, S. (2018). Pitt-Hopkins Syndrome: A Review of Current Literature, Clinical Approach, and 23-Patient Case Series. Journal of Child Neurology, 33(3), 233–244. 10.1177/0883073817750490

Hinderer, C., Bell, P., Vite, C. H., Louboutin, J.-P., Grant, R., Bote, E., Yu, H., Pukenas, B., Hurst, R., & Wilson, J. M. (2014). Widespread gene transfer in the central nervous system of cynomolgus macaques following delivery of AAV9 into the cisterna magna. Molecular Therapy. Methods & Clinical Development, 1, 14051. 10.1038/mtm.2014.51

Hordeaux, J., Wang, Q., Katz, N., Buza, E. L., Bell, P., & Wilson, J. M. (2018). The Neurotropic Properties of AAV-PHP.B Are Limited to C57BL/6J Mice. *Molecular Therapy*, *26*(3), 664–668. 10.1016/j.ymthe.2018.01.018

Huang, Q., Chan, K. Y., Wu, J., Botticello-Romero, N. R., Zheng, Q., Lou, S., Keyes, C., Svanbergsson, A., Johnston, J., Mills, A., Lin, C.-Y., Brauer, P. P., Clouse, G., Pacouret, S., Harvey, J. W., Beddow, T., Hurley, J. K., Tobey, I. G., Powell, M.,…Deverman, B. E. (2024). An AAV capsid reprogrammed to bind human transferrin receptor mediates brain-wide gene delivery. Science, 384(6701), 1220–1227. 10.1126/science.adm8386

Jung, M., Häberle, B. M., Tschaikowsky, T., Wittmann, M.-T., Balta, E.-A., Stadler, V.-C., Zweier, C., Dörfler, A., Gloeckner, C. J., & Lie, D. C. (2018). Analysis of the expression pattern of the schizophrenia-risk and intellectual disability gene TCF4 in the developing and adult brain suggests a role in development and plasticity of cortical and hippocampal neurons. Molecular Autism, 9, 20. 10.1186/s13229-018-0200-1

Kennedy, A. J., Rahn, E. J., Paulukaitis, B. S., Savell, K. E., Kordasiewicz, H. B., Wang, J., Lewis, J. W., Posey, J., Strange, S. K., Guzman-Karlsson, M. C., Phillips, S. E., Decker, K., Motley, S. T., Swayze, E. E., Ecker, D. J., Michael, T. P., Day, J. J., & Sweatt, J. D. (2016). Tcf4 regulates synaptic plasticity, DNA methylation, and memory function. Cell Reports, 16(10), 2666–2685. 10.1016/j.celrep.2016.08.004

Kernohan, K. D., & Boycott, K. M. (2024). The expanding diagnostic toolbox for rare genetic diseases. Nature Reviews. Genetics, 25(6), 401–415. 10.1038/s41576-023-00683-w

Kim, H., Berens, N. C., Ochandarena, N. E., & Philpot, B. D. (2020). Region and cell type distribution of TCF4 in the postnatal mouse brain. Frontiers in Neuroanatomy, 14, 42. 10.3389/fnana.2020.00042

Kim, H., Gao, E. B., Draper, A., Berens, N. C., Vihma, H., Zhang, X., Higashi-Howard, A., Ritola, K. D., Simon, J. M., Kennedy, A. J., & Philpot, B. D. (2022). Rescue of behavioral and electrophysiological phenotypes in a Pitt-Hopkins syndrome mouse model by genetic restoration of Tcf4 expression. ELife, 11. 10.7554/eLife.72290

Kronman, F. N., Liwang, J. K., Betty, R., Vanselow, D. J., Wu, Y.-T., Tustison, N. J., Bhandiwad, A., Manjila, S. B., Minteer, J. A., Shin, D., Lee, C. H., Patil, R., Duda, J. T., Xue, J., Lin, Y., Cheng, K. C., Puelles, L., Gee, J. C., Zhang, J.,…Kim, Y. (2024). Developmental mouse brain common coordinate framework. Nature Communications, 15(1), 9072. 10.1038/s41467-024-53254-w

Kügler, S., Kilic, E., & Bähr, M. (2003). Human synapsin 1 gene promoter confers highly neuron-specific long-term transgene expression from an adenoviral vector in the adult rat brain depending on the transduced area. Gene Therapy, 10(4), 337–347. 10.1038/sj.gt.3301905

Li, H., Zhu, Y., Morozov, Y. M., Chen, X., Page, S. C., Rannals, M. D., Maher, B. J., & Rakic, P. (2019). Disruption of TCF4 regulatory networks leads to abnormal cortical development and mental disabilities. Molecular Psychiatry, 24(8), 1235–1246. 10.1038/s41380-019-0353-0

Madisen, L., Zwingman, T. A., Sunkin, S. M., Oh, S. W., Zariwala, H. A., Gu, H., Ng, L. L., Palmiter, R. D., Hawrylycz, M. J., Jones, A. R., Lein, E. S., & Zeng, H. (2010). A robust and high-throughput Cre reporting and characterization system for the whole mouse brain. Nature Neuroscience, 13(1), 133–140. 10.1038/nn.2467

Marangi, G., Ricciardi, S., Orteschi, D., Lattante, S., Murdolo, M., Dallapiccola, B., Biscione, C., Lecce, R., Chiurazzi, P., Romano, C., Greco, D., Pettinato, R., Sorge, G., Pantaleoni, C., Alfei, E., Toldo, I., Magnani, C., Bonanni, P., Martinez, F.,…Zollino, M. (2011). The Pitt-Hopkins syndrome: report of 16 new patients and clinical diagnostic criteria. American Journal of Medical Genetics. Part A, 155A(7), 1536– 1545. 10.1002/ajmg.a.34070

Marangi, G., & Zollino, M. (2015). Pitt-Hopkins Syndrome and Differential Diagnosis: A Molecular and Clinical Challenge. Journal of Pediatric Genetics, 4(3), 168–176. 10.1055/s-0035-1564570

Mei, Y., Monteiro, P., Zhou, Y., Kim, J.-A., Gao, X., Fu, Z., & Feng, G. (2016). Adult restoration of Shank3 expression rescues selective autistic-like phenotypes. Nature, 530(7591), 481–484. 10.1038/nature16971

Mesman, S., Bakker, R., & Smidt, M. P. (2020). Tcf4 is required for correct brain development during embryogenesis. Molecular and Cellular Neurosciences, 106, 103502. 10.1016/j.mcn.2020.103502

Mielnik, C. A., Binko, M. A., Chen, Y., Funk, A. J., Johansson, E. M., Intson, K., Sivananthan, N., Islam, R., Milenkovic, M., Horsfall, W., Ross, R. A., Groc, L., Salahpour, A., McCullumsmith, R. E., Tripathy, S., Lambe, E. K., & Ramsey, A. J. (2021). Consequences of NMDA receptor deficiency can be rescued in the adult brain. Molecular Psychiatry, 26(7), 2929–2942. 10.1038/s41380-020-00859-4

Moser, E., Moser, M. B., & Andersen, P. (1993). Spatial learning impairment parallels the magnitude of dorsal hippocampal lesions, but is hardly present following ventral lesions. The Journal of Neuroscience, 13(9), 3916–3925. 10.1523/JNEUROSCI.13-09-03916.1993

Moser, M. B., Moser, E. I., Forrest, E., Andersen, P., & Morris, R. G. (1995). Spatial learning with a minislab in the dorsal hippocampus. Proceedings of the National Academy of Sciences of the United States of America, 92(21), 9697–9701. 10.1073/pnas.92.21.9697

Moyer, T. C., Hoffman, B. A., Chen, W., Shah, I., Ren, X.-Q., Knox, T., Liu, J., Wang, W., Li, J., Khalid, H., Kulkarni, A. S., Egbuchulam, M., Clement, J., Bloedel, A., Child, M., Kaur, R., Rouse, E., Graham, K., Maura, D.,…Nonnenmacher, M. E. (2025). Highly conserved brain vascular receptor ALPL mediates transport of engineered AAV vectors across the blood-brain barrier. Molecular Therapy, 33(8), 3902–3916. 10.1016/j.ymthe.2025.04.046

Nakazawa, K., McHugh, T. J., Wilson, M. A., & Tonegawa, S. (2004). NMDA receptors, place cells and hippocampal spatial memory. Nature Reviews. Neuroscience, 5(5), 361–372. 10.1038/nrn1385

Okamoto, K., Kamikubo, Y., Yamauchi, K., Okamoto, S., Takahashi, M., Ishida, Y., Koike, M., Ikegaya, Y., Sakurai, T., & Hioki, H. (2023). Specific AAV2/PHP.eB-mediated gene transduction of CA2 pyramidal cells via injection into the lateral ventricle. Scientific Reports, 13(1), 323. 10.1038/s41598-022-27372-8

Page, S. C., Hamersky, G. R., Gallo, R. A., Rannals, M. D., Calcaterra, N. E., Campbell, M. N., Mayfield, B., Briley, A., Phan, B. N., Jaffe, A. E., & Maher, B. J. (2018). The schizophrenia-and autism-associated gene, transcription factor 4 regulates the columnar distribution of layer 2/3 prefrontal pyramidal neurons in an activity-dependent manner. Molecular Psychiatry, 23(2), 304–315. 10.1038/mp.2017.37

Papes, F., Camargo, A. P., de Souza, J. S., Carvalho, V. M. A., Szeto, R. A., LaMontagne, E., Teixeira, J. R., Avansini, S. H., Sánchez-Sánchez, S. M., Nakahara, T. S., Santo, C. N., Wu, W., Yao, H., Araújo, B. M. P., Velho, P. E. N. F., Haddad, G. G., & Muotri, A. R. (2022). Transcription Factor 4 loss-of-function is associated with deficits in progenitor proliferation and cortical neuron content. Nature Communications, 13(1), 2387. 10.1038/s41467-022-29942-w

Phan, B. N., Bohlen, J. F., Davis, B. A., Ye, Z., Chen, H.-Y., Mayfield, B., Sripathy, S. R., Cerceo Page, S., Campbell, M. N., Smith, H. L., Gallop, D., Kim, H., Thaxton, C. L., Simon, J. M., Burke, E. E., Shin, J. H., Kennedy, A. J., Sweatt, J. D., Philpot, B. D.,…Maher, B. J. (2020). A myelin-related transcriptomic profile is shared by Pitt-Hopkins syndrome models and human autism spectrum disorder. Nature Neuroscience, 23(3), 375–385. 10.1038/s41593-019-0578-x

Rannals, M. D., Hamersky, G. R., Page, S. C., Campbell, M. N., Briley, A., Gallo, R. A., Phan, B. N., Hyde, T. M., Kleinman, J. E., Shin, J. H., Jaffe, A. E., Weinberger, D. R., & Maher, B. J. (2016). Psychiatric risk gene transcription factor 4 regulates intrinsic excitability of prefrontal neurons via repression of scn10a and KCNQ1. Neuron, 90(1), 43–55. 10.1016/j.neuron.2016.02.021

Sarkar, D., Shariq, M., Dwivedi, D., Krishnan, N., Naumann, R., Bhalla, U. S., & Ghosh, H. S. (2021). Adult brain neurons require continual expression of the schizophrenia-risk gene Tcf4 for structural and functional integrity. Translational Psychiatry, 11(1), 494. 10.1038/s41398-021-01618-x

Schaeren-Wiemers, N., Valenzuela, D. M., Frank, M., & Schwab, M. E. (1995). Characterization of a rat gene, rMAL, encoding a protein with four hydrophobic domains in central and peripheral myelin. The Journal of Neuroscience, 15(8), 5753– 5764. 10.1523/JNEUROSCI.15-08-05753.1995

Sepp, M., Kannike, K., Eesmaa, A., Urb, M., & Timmusk, T. (2011). Functional diversity of human basic helix-loop-helix transcription factor TCF4 isoforms generated by alternative 5’ exon usage and splicing. Plos One, 6(7), e22138. 10.1371/journal.pone.0022138

Silva-Santos, S., van Woerden, G. M., Bruinsma, C. F., Mientjes, E., Jolfaei, M. A., Distel, B., Kushner, S. A., & Elgersma, Y. (2015). Ube3a reinstatement identifies distinct developmental windows in a murine Angelman syndrome model. The Journal of Clinical Investigation, 125(5), 2069–2076. 10.1172/JCI80554

Sirp, A., Shubina, A., Tuvikene, J., Tamberg, L., Kiir, C. S., Kranich, L., & Timmusk, T. (2022). Expression of alternative transcription factor 4 mRNAs and protein isoforms in the developing and adult rodent and human tissues. Frontiers in Molecular Neuroscience, 15, 1033224. 10.3389/fnmol.2022.1033224

Thaxton, C., Kloth, A. D., Clark, E. P., Moy, S. S., Chitwood, R. A., & Philpot, B. D. (2018). Common Pathophysiology in Multiple Mouse Models of Pitt-Hopkins Syndrome. The Journal of Neuroscience, 38(4), 918–936. 10.1523/JNEUROSCI.1305-17.2017

Wallace, J. L., & Pollen, A. A. (2024). Human neuronal maturation comes of age: cellular mechanisms and species differences. Nature Reviews. Neuroscience, 25(1), 7–29. 10.1038/s41583-023-00760-3

Whalen, S., Héron, D., Gaillon, T., Moldovan, O., Rossi, M., Devillard, F., Giuliano, F., Soares, G., Mathieu-Dramard, M., Afenjar, A., Charles, P., Mignot, C., Burglen, L., Van Maldergem, L., Piard, J., Aftimos, S., Mancini, G., Dias, P., Philip, N.,…Giurgea, I. (2012). Novel comprehensive diagnostic strategy in Pitt-Hopkins syndrome: clinical score and further delineation of the TCF4 mutational spectrum. Human Mutation, 33(1), 64–72. 10.1002/humu.21639

Workman, A. D., Charvet, C. J., Clancy, B., Darlington, R. B., & Finlay, B. L. (2013). Modeling transformations of neurodevelopmental sequences across mammalian species. The Journal of Neuroscience, 33(17), 7368–7383. 10.1523/JNEUROSCI.5746-12.2013

Yardeni, T., Eckhaus, M., Morris, H. D., Huizing, M., & Hoogstraten-Miller, S. (2011). Retro-orbital injections in mice. Lab Animal, 40(5), 155–160. 10.1038/laban0511-155

Zalewska, A. (2019). Developmental milestones in neonatal and juvenile C57Bl/6 mouse - Indications for the design of juvenile toxicity studies. Reproductive Toxicology, 88, 91–128. 10.1016/j.reprotox.2019.07.019

Zhang, J., Wu, Y., Chen, S., Luo, Q., Xi, H., Li, J., Qin, X., Peng, Y., Ma, N., Yang, B., Qiu, X., Lu, W., Chen, Y., Jiang, Y., Chen, P., Liu, Y., Zhang, C., Zhang, Z., Xiong, Y.,…Huang, H. (2024). Prospective prenatal cell-free DNA screening for genetic conditions of heterogenous etiologies. Nature Medicine, 30(2), 470–479. 10.1038/s41591-023-02774-x

Zollino, M., Zweier, C., Van Balkom, I. D., Sweetser, D. A., Alaimo, J., Bijlsma, E. K., Cody, J., Elsea, S. H., Giurgea, I., Macchiaiolo, M., Smigiel, R., Thibert, R. L., Benoist, I., Clayton-Smith, J., De Winter, C. F., Deckers, S., Gandhi, A., Huisman, S., Kempink, D.,…Hennekam, R. C. (2019). Diagnosis and management in Pitt-Hopkins syndrome: First international consensus statement. Clinical Genetics, 95(4), 462–478. 10.1111/cge.13506

Zweier, C, Sticht, H., Bijlsma, E. K., Clayton-Smith, J., Boonen, S. E., Fryer, A., Greally, M. T., Hoffmann, L., den Hollander, N. S., Jongmans, M., Kant, S. G., King, M. D., Lynch, S. A., McKee, S., Midro, A. T., Park, S. M., Ricotti, V., Tarantino, E., Wessels, M.,…Rauch, A. (2008). Further delineation of Pitt-Hopkins syndrome: phenotypic and genotypic description of 16 novel patients. Journal of Medical Genetics, 45(11), 738–744. 10.1136/jmg.2008.060129

Zweier, Christiane, Peippo, M. M., Hoyer, J., Sousa, S., Bottani, A., Clayton-Smith, J., Reardon, W., Saraiva, J., Cabral, A., Gohring, I., Devriendt, K., de Ravel, T., Bijlsma, E. K., Hennekam, R. C. M., Orrico, A., Cohen, M., Dreweke, A., Reis, A., Nurnberg, P., & Rauch, A. (2007). Haploinsufficiency of TCF4 causes syndromal mental retardation with intermittent hyperventilation (Pitt-Hopkins syndrome). American Journal of Human Genetics, 80(5), 994–1001. 10.1086/515583

